# Predicting commercially available antiviral drugs that may act on the novel coronavirus (2019-nCoV), Wuhan, China through a drug-target interaction deep learning model

**DOI:** 10.1101/2020.01.31.929547

**Authors:** Bo Ram Beck, Bonggun Shin, Yoonjung Choi, Sungsoo Park, Keunsoo Kang

**Affiliations:** Deargen, Inc., Daejeon, Republic of Korea; Department of Computer Science, Emory University, Atlanta, GA, United States; Department of Microbiology, College of Natural Sciences, Dankook University, Cheonan, Republic of Korea

## Abstract

The infection of a novel coronavirus found in Wuhan of China (2019-nCoV) is rapidly spreading, and the incidence rate is increasing worldwide. Due to the lack of effective treatment options for 2019-nCoV, various strategies are being tested in China, including drug repurposing. In this study, we used our pretrained deep learning-based drug-target interaction model called Molecule Transformer-Drug Target Interaction (MT-DTI) to identify commercially available drugs that could act on viral proteins of 2019-nCoV. The result showed that atazanavir, an antiretroviral medication used to treat and prevent the human immunodeficiency virus (HIV), is the best chemical compound, showing a inhibitory potency with *K_d_* of 94.94 nM against the 2019-nCoV 3C-like proteinase, followed by efavirenz (199.17 nM), ritonavir (204.05 nM), and dolutegravir (336.91 nM). Interestingly, lopinavir, ritonavir, and darunavir are all designed to target viral proteinases. However, in our prediction, they may also bind to the replication complex components of 2019-nCoV with an inhibitory potency with *K_d_* < 1000 nM. In addition, we also found that several antiviral agents, such as Kaletra, could be used for the treatment of 2019-nCoV, although there is no real-world evidence supporting the prediction. Overall, we suggest that the list of antiviral drugs identified by the MT-DTI model should be considered, when establishing effective treatment strategies for 2019-nCoV.

## Introduction

Coronaviruses (CoVs), belonging to the family *Coronaviridae*, are positive-sense enveloped RNA viruses and cause infections in birds, mammals, and humans (1–3). The family includes four genera, such as *Alphacoronavirus*, *Betacoronavirus*, *Deltacoronavirus*, and *Gammacoronavirus* (4). Two infamous infectious coronaviruses in the genus *Betacoronavirus* are severe acute respiratory syndrome coronavirus (SARS-CoV) (5) and Middle East respiratory syndrome coronavirus (MERS-CoV) (6), which have infected more than 10,000 people around the world in the past two decades. Unfortunately, the incidence was accompanied by high mortality rates (9.6% for SARS-CoV and 34.4% for MERS-CoV), indicating that there is an urgent need for effective treatment at the beginning of the outbreak to prevent the spread (7, 8). However, this cannot be achieved with current drug development or an application system, taking several years for newly developed drugs to come to the market. Unexpectedly, the world is facing the same situation as the previous outbreak due to a recent epidemic of atypical pneumonia caused by a novel coronavirus (2019-nCoV) in Wuhan, China (5, 9). Thus, a rapid drug application strategy that can be immediately applied to the patient is necessary. Currently, the only way to address this matter is to repurpose commercially available drugs for the pathogen in so-called “drug-repurposing”. However, in theory, artificial intelligence (AI)-based architectures must be taken into account in order to accurately predict drug-target interactions (DTIs). This is because of the enormous amount of complex information (e.g. hydrophobic interactions, ionic interactions, hydrogen bonding, and/or van der Waals forces) between molecules. To this end, we previously developed a deep learning-based drug-target interaction prediction model, called Molecule Transformer-Drug Target Interaction (MT-DTI) (10).

In this study, we applied our pre-trained MT-DTI model to identify commercially available antiviral drugs that could potentially disrupt 2019-nCoV’s viral components, such as proteinase, RNA-dependent RNA polymerase, and/or helicase. Since the model utilizes simplified molecular-input line-entry system (SMILES) strings and amino acid (AA) sequences, which are 1D string inputs, it is possible to quickly apply target proteins that do not have experimentally confirmed 3D crystal structures, such as viral proteins of 2019-nCoV. We share a list of top commercially available antiviral drugs that could potentially hinder the multiplication cycle of 2019-nCoV with the hope that effective drugs can be developed based on these AI-proposed drug candidates and act against 2019-nCoV.

## Methods

### Amino acid sequences used in this study

Amino acid sequences of 3C-like proteinase (accession YP_009725301.1), RNA-dependent RNA polymerase (accession YP_009725307.1), helicase (accession YP_009725308.1), 3’-to-5’ exonuclease (accession YP_009725309.1), endoRNAse (accession YP_009725310.1), and 2’-O-ribose methyltransferase (accession YP_009725311.1) of the 2019-nCoV replication complex were extracted from the 2019-nCoV whole genome sequence (accession NC_045512.2), from the National Center for Biotechnology Information (NCBI) database. The raw prediction results were screened for antiviral drugs that are FDA approved, target viral proteins, and have a *K_d_* value less than 1,000 nM.

### Prediction of drug-target interactions using binding affinity scores

Molecule transformer-drug target interaction (MT-DTI) was used to predict binding affinity values between commercially available antiviral drugs and target proteins. Briefly, the natural language processing (NLP) based Bidirectional Encoder Representations from Transformers (BERT) framework is a core algorithm of the model with good performance and robust results in diverse drug-target interaction datasets through pretraining with ‘chemical language’ SMILES of approximately 1,000,000,000 compounds.

To train the model, the Drug Target Common (DTC) database (11) and BindingDB (12) database were manually curated and combined. Three types of efficacy value, *K_i_*, *K_d_*, and IC_50_ were integrated by a consistence-score-based averaging algorithm (13) to make the Pearson correlation score over 0.9 in terms of *K_i_*, *K_d_*, and IC_50_. Since the BindingDB database includes a wide variety of species and target proteins, the MT-DTI model has the potential power to predict interactions between antiviral drugs and 2019-nCoV proteins.

## Results and Discussion

The 2019-nCoV 3C-like proteinase was predicted to bind with atazanavir (*K_d_* 94.94 nM), followed by efavirenz, ritonavir, and other antiviral drugs that have a predicted affinity of *K_d_* > 100 nM potency (Table 1). No other protease inhibitor antiviral drug was found in the *K_d_* < 1,000 nM range. Although there is no real-world evidence about whether these drugs will act as predicted against 2019-nCoV yet, some case studies have been identified. For example, a docking study of lopinavir along with other HIV proteinase inhibitors of the CoV proteinase (PDBID 1UK3) suggests atazanavir and ritonavir, which are listed in the present prediction results, may inhibit the CoV proteinase in line with the inhibitory potency of lopinavir (14). According to the prediction, viral proteinase-targeting drugs were predicted to act more favorably on the viral replication process than viral proteinase through the DTI model (Table 2–6). The results include antiviral drugs other than proteinase inhibitors, such as guanosine analogues (e.g., acyclovir, ganciclovir, and penciclovir), reverse transcriptase inhibitors, and integrase inhibitors.

**Table 1.** Drug-target interaction (DTI) prediction results of FDA approved antviral drugs available on markets against a novel coronavirus (2019-nCoV, NCBI reference sequence NC_045512.2) 3C-like proteinase (accession YP_009725301.1). Ritonavir is expressed in canonical and isomeric forms SMILES, and * indicates isomeric form SMILES of ritonavir.

| Small molecules | SMILES used in the DTI prediction | $K_d$ in nM |
| --- | --- | --- |
| Atazanavir | COC(=O)NC(C(=O)NC(Cc1ccccc1)C(O)CN(Cc1ccc(-c2ccccc2)cc1)NC(=O)C(NC(=O)OC)C(C)(C)C(C)(C)C | 94.94 |
| Efavirenz | O=C1Nc2ccc(Cl)cc2[C@@](C#CC2CC2)(C(F)(F)F)O1 | 199.17 |
| Ritonavir | CC(C)c1nc(CN(C)C(=O)NC(C(=O)NC(Cc2ccccc2)CC(O)C(Cc2ccccc2)NC(=O)OCc2cncc2)C(C)C)cs1 | 204.05 |
| Dolutegravir | CC1CCOC2Cn3cc(C(=O)NCc4ccc(F)cc4F)c(=O)c(O)c3C(=O)N12 | 336.91 |
| Asunaprevir | C=CC1CC1(NC(=O)C1CC(Oc2ncc(OC)c3ccc(Cl)cc23)CN1C(=O)C(NC(=O)OC(C)(C)C(C)(C)C(C)C(=O)NS(=O)(=O)C1CC1 | 581.77 |
| Ritonavir* | CC(C)c1nc(CN(C)C(=O)N[C@H](C(=O)N[C@@H](Cc2ccccc2)C[C@H](O)[C@H](Cc2ccccc2)NC(=O)OCc2cncc2)C(C)C)cs1 | 609.02 |
| Simeprevir | COc1ccc2c(O[C@H]3CC4C(=O)N(C)CCCC/C=C/[C@H]5C[C@@]5(C(=O)NS(=O)(=O)C5CC5)NC(=O)[C@@H]4C3)cc(-c3nc(C(C)C)cs3)nc2c1C | 826.24 |

**Table 2.** Drug-target interaction (DTI) prediction results of antiviral drugs available on markets against a novel coronavirus (2019-nCoV, NCBI reference sequence NC_045512.2) RNA-dependent RNA polymerase (accession YP_009725307.1).

| Small molecules | SMILES used in the DTI prediction | $K_d$ in nM |
| --- | --- | --- |
| Grazoprevir | C=C[C@H]1C[C@]1(NC(=O)[C@@H]1C[C@@H]2CN1C(=O)[C@H](C(C)(C)C)NC(=O)O[C@@H]1C[C@H]1CCCCCc1nc3ccc(OC)cc3nc1O2)C(=O)NS(=O)(=O)C1CC1 | 8.69 |
| Ganciclovir | Nc1nc(=O)c2ncn(COC(CO)CO)c2[nH]1 | 11.91 |
| Atazanavir | COC(=O)NC(C(=O)NC(Cc1ccccc1)C(O)CN(Cc1ccc(-c2ccccc2)cc1)NC(=O)C(NC(=O)OC)C(C)(C)C(C)(C)C | 21.83 |
| Daclatasvir | COC(=O)NC(C(=O)N1CCCC1c1ncc(-c2ccc(-c3ccc(-c4cnc(C5CCCN5C(=O)C(NC(=O)OC)C(C)C)[nH]4)cc3)cc2)[nH]1)C(C)C | 23.31 |
| Acyclovir | Nc1nc(=O)c2ncn(COCCO)c2[nH]1 | 26.66 |
| Etravirine | Cc1cc(C#N)cc(C)c1Oc1nc(Nc2ccc(C#N)cc2)nc(N)c1Br | 33.09 |
| Entecavir | C=C1[C@@H](n2cnc3c(=O)nc(N)[nH]c32)C[C@H](O)[C@H]1CO | 52.83 |
| Efavirenz | O=C1Nc2ccc(Cl)cc2[C@@](C#CC2CC2)(C(F)(F)F)O1 | 76.70 |
| Asunaprevir | C=CC1CC1(NC(=O)C1CC(Oc2ncc(OC)c3ccc(Cl)cc23)CN1C(=O)C(NC(=O)OC(C)(C)C(C)(C)C(C)C(=O)NS(=O)(=O)C1CC1 | 78.36 |
| Abacavir | Nc1nc(NC2CC2)c2ncn(C3C=CC(CO)C3)c2n1 | 131.51 |
| Dolutegravir | CC1CCOC2Cn3cc(C(=O)NCc4ccc(F)cc4F)c(=O)c(O)c3C(=O)N12 | 150.15 |
| Lomibuvir | CC1CCC(C(=O)N(c2cc(C#CC(C)(C)C)sc2C(=O)O)C2CCC(O)CC2)CC1 | 280.96 |
| Penciclovir | Nc1nc(=O)c2ncn(CCC(CO)CO)c2[nH]1 | 312.93 |
| Trifluridine | O=c1[nH]c(=O)n(C2CC(O)C(CO)O2)cc1C(F)(F)F | 315.79 |
| Danoprevir | CC(C)(C)OC(=O)N[C@@H]1CCCCC/C=C/[C@@H]2C[C@@]2(CNS(=O)(=O)C2CC2)NC(=O)[C@@H]2C[C@@H](OC(=O)N3Cc4cccc(F)c4C3)CN2C1=O | 405.66 |
| Ritonavir | CC(C)c1nc(CN(C)C(=O)NC(C(=O)NC(Cc2ccccc2)CC(O)C(Cc2ccccc2)NC(=O)OCc2cncs2)C(C)C)cs1 | 624.30 |
| Saquinavir | CC(C)(C)NC(=O)[C@@H]1C[C@@H]2CCCC[C@H]2CN1C[C@@H](O)[C@H](Cc1ccccc1)NC(=O)[C@H](CC(N)=O)NC(=O)c1ccc2ccccc2n1 | 704.86 |
| Raltegravir | Cc1nnc(C(=O)NC(C)(C)c2nc(C(=O)NCc3ccc(F)cc3)c(O)c(=O)n2C)o1 | 832.25 |
| Lamivudine | Nc1ccn([C@@H]2CS[C@H](CO)O2)c(=O)n1 | 999.92 |

Among the prediction results, atazanavir was predicted to have a potential binding affinity to bind to RNA-dependent RNA polymerase (*K_d_* 21.83 nM), helicase (*K_d_* 25.92 nM), 3’-to-5’ exonuclease (*K_d_* 82.36 nM), 2’-O-ribose methyltransferase (*K_d_* of 390 nM), and endoRNAse (*K_d_* 50.32 nM), which suggests that all subunits of the 2019-nCoV replication complex may be inhibited simultaneously by atazanavir (Table 2–6). Also, ganciclovir was predicted to bind to three subunits of the replication complex of the 2019-nCoV: RNA-dependent RNA polymerase (*K_d_* 11.91 nM), 3’-to-5’ exonuclease (*K_d_* 56.29 nM), and RNA helicase (*K_d_* 108.21 nM). Lopinavir and ritonavir, active materials of AbbVie’s Kaletra, both were predicted to have a potential affinity to 2019-nCoV helicase (Table 3) and are suggested as potential MERS therapeutics (15). Recently, approximately $2 million worth of Kaletra doses were donated to China (16), and a previous clinical study of SARS by Chu et al. (17) may support this decision (17). Another anti-HIV drug, Prezcobix of Johnson & Johnson, which consists of darunavir and cobicistat, was to be sent to China (16), and darunavir is also predicted to have a *K_d_* of 90.38 nM against 2019-nCoV’s helicase (Table 3). However, there was no current supporting literature found for darunavir to be used as a CoV therapeutic.

**Table 3.** Drug-target interaction (DTI) prediction results of antiviral drugs available on markets against a novel coronavirus (2019-nCoV, NCBI reference sequence NC_045512.2) helicase (accession YP_009725308.1). Ritonavir is expressed in canonical and isomeric form SMILES, and * indicates isomeric form SMILES of ritonavir.

| Small molecules | SMILES used in the DTI prediction | K<sub>d</sub> in nM |
| --- | --- | --- |
| Simeprevir | COc1ccc2c(O[C@H]3CC4C(=O)N(C)CCCC/C=C/[C@H]5C[C@@]5(C(=O)NS(=O)(=O)C5CC5)NC(=O)[C@@H]4C3)cc(-c3nc(C(C)C)cs3)nc2c1C | 23.34 |
| Atazanavir | COC(=O)NC(C(=O)NC(Cc1ccccc1)C(O)CN(Cc1ccc(-c2ccccc2)cc1)NC(=O)C(NC(=O)OC)C(C)(C)C)C(C)C | 25.92 |
| Grazoprevir | C=C[C@H]1C[C@]1(NC(=O)[C@@H]1C[C@@H]2CN1C(=O)[C@H](C(C)(C)C)NC(=O)O[C@@H]1C[C@H]1CCCCc1nc3ccc(OC)cc3nc1O2)C(=O)NS(=O)(=O)C1CC1 | 26.28 |
| Asunaprevir | C=CC1CC1(NC(=O)C1CC(OC2ncc(OC)c3ccc(Cl)cc23)CN1C(=O)C(NC(=O)OC(C)(C)C)C(C)C)C(=O)NS(=O)(=O)C1CC1 | 28.20 |
| Telaprevir | CCCC(NC(=O)C1C2CCCC2CN1C(=O)C(NC(=O)C(NC(=O)c1nccn1)C1CCCCC1)C(C)(C)C)C(=O)C(=O)NC1CC1 | 40.75 |
| Ritonavir | CC(C)c1nc(CN(C)C(=O)NC(C(=O)NC(Cc2ccccc2)CC(O)C(Cc2ccccc2)NC(=O)OCc2cncs2)C(C)C)cs1 | 41.60 |
| Lopinavir | Cc1cccc(C)c1OCC(=O)N[C@@H](Cc1ccccc1)[C@@H](O)C[C@H](Cc1ccccc1)NC(=O)[C@H](C(C)C)N1CCCN1=O | 78.49 |
| Darunavir | CC(C)CN(C[C@@H](O)[C@H](Cc1ccccc1)/N=C(\O)O[C@H]1C O[C@H]2OCC[C@@H]12)S(=O)(=O)c1ccc(N)cc1 | 90.38 |
| Ganciclovir | Nc1nc(=O)c2ncn(COC(CO)CO)c2[nH]1 | 108.21 |
| Penciclovir | Nc1nc(=O)c2ncn(CCC(CO)CO)c2[nH]1 | 129.41 |
| Etravirine | Cc1cc(C#N)cc(C)c1Oc1nc(Nc2ccc(C#N)cc2)nc(N)c1Br | 175.50 |
| Raltegravir | Cc1nnc(C(=O)NC(C)(C)c2nc(C(=O)NCc3ccc(F)cc3)c(O)c(=O)n2C)o1 | 299.81 |
| Dolutegravir | CC1CCOC2Cn3cc(C(=O)NCc4ccc(F)cc4F)c(=O)c(O)c3C(=O)N12 | 333.32 |
| Nelfinavir | Cc1c(O)cccc1C(=O)NC(CSc1cccc1)C(O)CN1CC2CCCCC2CC1C(=O)NC(C)(C)C | 365.96 |
| Indinavir | CC(C)(C)NC(=O)[C@@H]1CN(Cc2cccnc2)CCN1C[C@@H](O)C[C@@H](Cc1cccc1)C(=O)N[C@H]1c2cccc2C[C@H]1O | 401.78 |
| Efavirenz | O=C1Nc2ccc(Cl)cc2C(C#CC2CC2)(C(F)(F)F)O1 | 412.86 |
| Entecavir | C=C1[C@@H](n2cnc3c(=O)nc(N)[nH]c32)C[C@H](O)[C@H]1CO | 452.78 |
| Ritonavir* | CC(C)c1nc(CN(C)C(=O)N[C@H](C(=O)N[C@@H](Cc2cccc2)C[C@H](O)[C@H](Cc2cccc2)NC(=O)OCc2cncs2)C(C)C)cs1 | 462.20 |
| Boceprevir | CC(C)(C)NC(=O)NC(C(=O)N1CC2C(C1C(=O)NC(CC1CCC1)C(=O)C(N)=O)C2(C)C)C(C)(C)C | 510.35 |
| Lomibuvir | CC1CCC(C(=O)N(c2cc(C#CC(C)(C)C)sc2C(=O)O)C2CCC(O)C2)CC1 | 543.41 |
| Acyclovir | Nc1nc(=O)c2ncn(COCCO)c2[nH]1 | 661.76 |

**Table 4.** Drug-target interaction (DTI) prediction results of anti-viral drugs available on markets against a novel coronavirus (2019-nCoV, NCBI reference sequence NC_045512.2) 3’-to-5’ exonuclease (accession YP_009725309.1).

| Small molecules | SMILES used in the DTI prediction | $K_d$ in nM |
| --- | --- | --- |
| Simeprevir | COc1ccc2c(O[C@H]3CC4C(=O)N(C)CCCC/C=C/[C@H]5C[C@@]5(C(=O)NS(=O)(=O)C5CC5)NC(=O)[C@@H]4C3)cc(-c3nc(C(C)C)cs3)nc2c1C | 13.40 |
| Efavirenz | O=C1Nc2ccc(Cl)cc2[C@@H](C#CC2CC2)(C(F)(F)F)O1 | 39.55 |
| Danoprevir | CC(C)(C)OC(=O)N[C@H]1CCCC/C=C/[C@@H]2C[C@@]2(CNS(=O)(=O)C2CC2)NC(=O)[C@@H]2C[C@@H](OC(=O)N3Cc4cccc(F)c4C3)CN2C1=O | 49.26 |
| Ganciclovir | Nc1nc(=O)c2ncn(COC(CO)CO)c2[nH]1 | 56.29 |
| Penciclovir | Nc1nc(=O)c2ncn(CCC(CO)CO)c2[nH]1 | 71.76 |
| Atazanavir | COC(=O)NC(C(=O)NC(Cc1cccc1)C(O)CN(Cc1ccc(-c2ccccn2)cc1)NC(=O)C(NC(=O)OC)C(C)(C)C)C(C)(C)C | 82.36 |
| Entecavir | C=C1[C@@H](n2cnc3c(=O)nc(N)[nH]c32)C[C@H](O)[C@H]1CO | 82.78 |
| Daclatasvir | COC(=O)NC(C(=O)N1CCCC1c1ncc(-c2ccc(-c3ccc(-c4cnc(C5CCCN5C(=O)C(NC(=O)OC)C(C)C)[nH]4)cc3)cc2)[nH]1)C(C)C | 110.47 |
| Grazoprevir | C=C[C@H]1C[C@]1(NC(=O)[C@@H]1C[C@@H]2CN1C(=O)[C@H](C(C)(C)C)NC(=O)O[C@@H]1C[C@H]1CCCCc1nc3ccc(OC)cc3nc1O2)C(=O)NS(=O)(=O)C1CC1 | 111.90 |
| Asunaprevir | C=CC1CC1(NC(=O)C1CC(OC2ncc(OC)c3ccc(Cl)cc23)CN1C(=O)C(NC(=O)OC(C)(C)C)C(C)(C)C(=O)NS(=O)(=O)C1CC1 | 117.26 |
| Ritonavir | CC(C)c1nc(CN(C)C(=O)NC(C(=O)NC(Cc2cccc2)CC(O)C(Cc2cccc2)NC(=O)OCc2cncs2)C(C)C)cs1 | 182.51 |
| Lomibuvir | CC1CCC(C(=O)N(c2cc(C#CC(C)(C)C)sc2C(=O)O)C2CCC(O)C2)CC1 | 182.65 |
| Darunavir | CC(C)CN[C@@H](O)[C@H](Cc1cccc1)/N=C(\O)O[C@H]1CO[C@H]2OCC[C@@H]12S(=O)(=O)c1ccc(N)cc1 | 195.73 |
| Raltegravir | Cc1nnc(C(=O)NC(C)(C)c2nc(C(=O)NCc3ccc(F)cc3)c(O)c(=O)n2C)o1 | 306.99 |
| Dolutegravir | CC1CCOC2Cn3cc(C(=O)NCc4ccc(F)cc4F)c(=O)c(O)c3C(=O)N12 | 326.89 |
| Lopinavir | Cc1cccc(C)c1OCC(=O)NC(Cc1cccc1)C(O)CC(Cc1cccc1)NC(=O)C(C(C)C)N1CCCNC1=O | 959.76 |

**Table 5.** Drug-target interaction (DTI) prediction results of anti-viral drugs available on markets against a novel coronavirus (2019-nCoV, NCBI reference sequence NC_045512.2) endoRNAse (accession YP_009725310.1).

| Small molecules | SMILES used in the DTI prediction | $K_d$ in nM |
| --- | --- | --- |
| Efavirenz | O=C1Nc2ccc(Cl)cc2[C@@](C#CC2CC2)(C(F)(F)F)O1 | 34.19 |
| Atazanavir | COC(=O)NC(C(=O)NC(Cc1cccc1)C(O)CN(Cc1ccc(-c2ccccc2)cc1)NC(=O)C(NC(=O)OC)C(C)(C)C(C)(C)C | 50.32 |
| Ritonavir | CC(C)c1nc(CN(C)C(=O)NC(C(=O)NC(Cc2ccccc2)CC(O)C(Cc2ccccc2)NC(=O)OCc2cncs2)C(C)C)cs1 | 124.36 |
| Danoprevir | CC(C)(C)OC(=O)N[C@H]1CCCC/C=C/[C@@H]2C[C@@]2(CNS(=O)(=O)C2CC2)NC(=O)[C@@H]2C[C@@H](OC(=O)N3Cc4cccc(F)c4C3)CN2C1=O | 235.15 |
| Grazoprevir | C=C[C@H]1C[C@]1(NC(=O)[C@@H]1C[C@@H]2CN1C(=O)[C@H](C(C)(C)C)NC(=O)O[C@@H]1C[C@H]1CCCCc1nc3ccc(OC)cc3nc1O2)C(=O)NS(=O)(=O)C1CC1 | 277.87 |
| Dolutegravir | CC1CCOC2Cn3cc(C(=O)NCc4ccc(F)cc4F)c(=O)c(O)c3C(=O)N12 | 349.63 |
| Lomibuvir | CC1CCC(C(=O)N(c2cc(C#CC(C)(C)C)sc2C(=O)O)C2CCC(O)C2)CC1 | 398.81 |
| Lopinavir | Cc1cccc(C)c1OCC(=O)N[C@@H](Cc1cccc1)[C@@H](O)C[C@@H](Cc1cccc1)NC(=O)[C@H](C(C)C)N1CCCNC1=O | 472.08 |
| Darunavir | CC(C)CN(C[C@@H](O)[C@H](Cc1cccc1)/N=C(\O)O[C@H]1CO[C@H]2OCC[C@@H]12)S(=O)(=O)c1ccc(N)cc1 | 562.40 |
| Nelfinavir | Cc1c(O)cccc1C(=O)NC(CSc1cccc1)C(O)CN1CC2CCCCC2CC1C(=O)NC(C)(C)C | 576.82 |
| Telaprevir | CCCC(NC(=O)C1C2CCCC2CN1C(=O)C(NC(=O)C(NC(=O)c1nccn1)C1CCCC1)C(C)(C)C(=O)C(=O)NC1CC1 | 618.11 |
| Abacavir | Nc1nc(NC2CC2)c2ncn(C3C=CC(CO)C3)c2n1 | 619.79 |
| Raltegravir | Cc1nnc(C(=O)NC(C)(C)c2nc(C(=O)NCc3ccc(F)cc3)c(O)c(=O)n2C)o1 | 727.37 |
| Boceprevir | CC(C)(C)NC(=O)NC(C(=O)N1CC2C(C1C(=O)NC(CC1CCC1)C(=O)C(N)=O)C2(C)C)C(C)(C)C | 891.62 |

**Table 6.** Drug-target interaction (DTI) prediction results of anti-viral drugs available on markets against a novel coronavirus (2019-nCoV, NCBI reference sequence NC_045512.2) 2’-O-ribose methyltransferase (accession YP_009725311.1).

| Small molecules | SMILES used in the DTI prediction | $K_d$ in nM |
| --- | --- | --- |
| Atazanavir | COC(=O)NC(C(=O)NC(Cc1ccccc1)C(O)CN(Cc1ccc(-c2ccccc2)cc1)NC(=O)C(NC(=O)OC)C(C)(C)C(C)(C)C | 390.67 |
| Efavirenz | O=C1Nc2ccc(Cl)cc2[C@@](C#CC2CC2)(C(F)(F)F)O1 | 423.00 |
| Boceprevir | CC(C)(C)NC(=O)NC(C(=O)N1CC2C(C1C(=O)NC(CC1CCC1)C(=O)C(N)=O)C2(C)C)C(C)(C)C | 433.93 |

## Conclusion

In many cases, DTI prediction models serve as a tool to repurpose drugs to develop novel usages of existing drugs. However, the application of DTI prediction in the present study may be useful to control unexpected and rapidly spreading infections such SARS-CoV, MERS-CoV, and 2019-nCoV at the frontline of the disease control until better therapeutic measures are developed. Nevertheless, we hope our prediction results may support and helpful to experimental therapy options for China and other countries identified 2019-nCoV infection.

